# Direction-dependent misalignment between joint and muscle spaces predicts different generalization patterns following adaptation to opposing visuomotor rotations

**DOI:** 10.64898/2026.07.20.739526

**Authors:** Luca Creitz, Sergio Gurgone, Ryosuke Murai, Nobuhiro Hagura, Johannes Maria Nicolaas Essers, Tsuyoshi Ikegami

**Affiliations:** Center for Information and Neural Networks (CiNet), Advanced ICT Research Institute, National Institute of Information and Communications Technology, 1-4, Yamadaoka, Suita City, Osaka, Japan; Faculty of Health, Medicine and Life Sciences, Maastricht University, Maastricht, the Netherlands; Institute NUTRIM for Nutrition and Translational Research in Metabolism, Maastricht, the Netherlands; Graduate School of Frontier Biosciences, Osaka University, 1-3, Yamadaoka, Suita, Osaka, Japan

**Keywords:** : Muscle space, Joint space, Motor representation, Motor generalization, Visuomotor adaptation

## Abstract

Learned movements are thought to be represented in both extrinsic and intrinsic reference frames. Intrinsic representations are traditionally characterized using joint-based coordinates, although the relationship between muscle activation patterns and joint movements strongly depends on limb configuration. Consequently, movement directions aligned in joint space may not be aligned in muscle space. The implications of this mismatch for motor learning, however, have remained largely unexplored. We addressed this question by combining muscle-pattern similarity analysis across workspaces with generalization analysis of a visuomotor adaptation task. In Experiment 1, participants performed planar reaching movements in two workspaces separated by a 45° shoulder rotation while electromyographic activity was recorded from nine upper-limb muscles. Similarity analysis revealed that movement directions aligned in joint space between the two postures were not always aligned in muscle space and that the degree of misalignment varied with movement direction. Based on these findings, we predicted that visuomotor adaptation to clockwise (CW) and counterclockwise (CCW) rotations would produce different patterns of motor generalization, in contrast to the identical generalization patterns predicted by a joint-space account. Experiment 2 confirmed this prediction, revealing a systematic shift between the CW and CCW generalization patterns that was consistent with the prediction based on muscle-pattern similarity. These findings suggest that intrinsic representations of learned movements cannot be fully captured by joint-based coordinates alone and that muscle-based coordinates contribute to the representation and generalization of learned movements. Together, these findings highlight the importance of considering the underlying biomechanics when interpreting motor representations using generalization paradigms.

## Introduction

The brain can acquire and retain highly complex motor skills, from playing the piano to dancing intricate choreographies, and with sufficient practice these skills are performed with little conscious mental effort. Yet how learned movements are represented in the brain remains a central question in neuroscience. A fundamental aspect of this question concerns the reference frames in which learned movements are represented. Understanding these reference frames would therefore provide key insights into the computational organization of motor learning (Shadmehr 2004).

Previous research has proposed that learned movements are represented in two classes of reference frames: extrinsic (Krakauer et al., 2000, Vetter et al., 1999, Wang and Sainburg, 2005) and intrinsic (Malfait et al., 2002, Shadmehr and Moussavi, 2000, Shadmehr and Mussa-Ivaldi, 1994). Extrinsic representations describe movements in Cartesian coordinates, such as changes in hand position (Fig. 1A). In contrast, the intrinsic representations describe movements in body-based coordinates, such as changes in joint angles (Fig. 1B). To examine how learned movements are represented across these reference frames, prior studies have employed motor generalization paradigms. In these paradigms, participants adapt to a perturbation (either force fields or visuomotor rotations) while reaching to a training target, after which generalization is examined in an untrained workspace with different arm postures and movement directions (Shadmehr and Mussa-Ivaldi, 1994, Shadmehr and Wise, 2005, Brayanov et al., 2012, Poh and Taylor, 2019). Depending on the coordinate system in which the learned movement is represented, different generalization patterns can be predicted.

**Figure 1:**
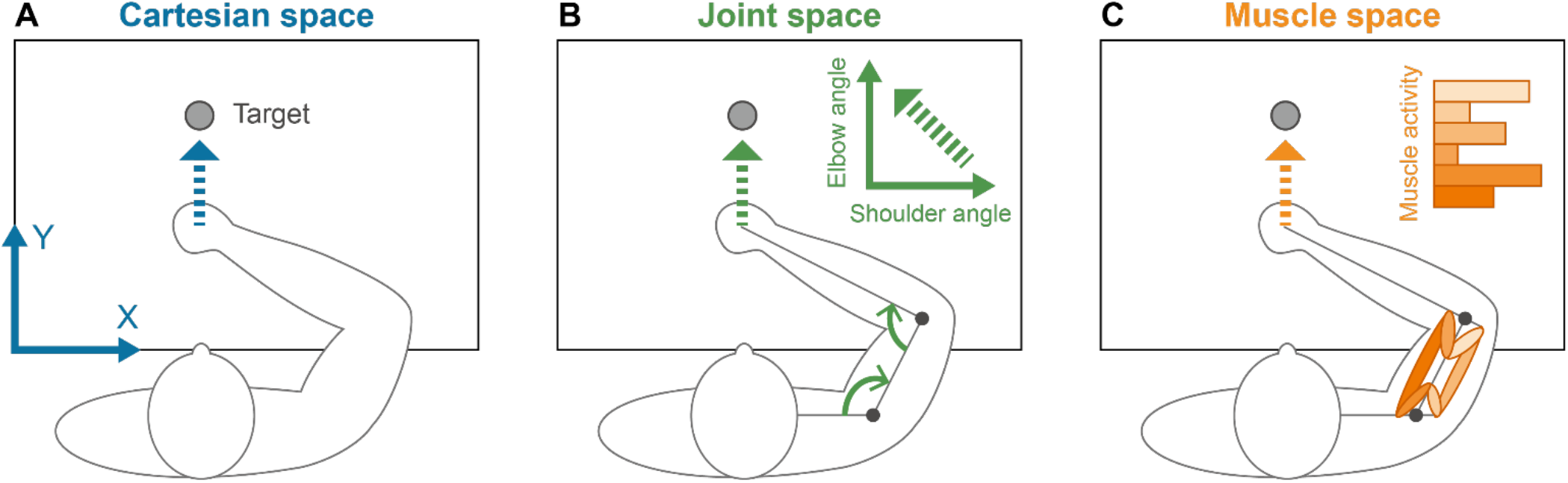
Schematic illustration of three spaces for movement representation. (A) Cartesian space, in which movements are represented as changes in the end-effector (hand) position. (B) Joint space, in which movements are represented as changes in joint angles. (C) Muscle space, in which movements are represented by muscle activity patterns.

Previous studies have reported conflicting findings: force-field adaptation studies have supported intrinsic representations (Haswell et al., 2009, Malfait et al., 2002, Shadmehr and Mussa-Ivaldi, 1994), whereas visuomotor adaptation studies have supported extrinsic representations (Krakauer et al., 2000, Wang and Sainburg, 2005, Poh et al., 2017, Franklin et al., 2016, Malfait et al., 2005). However, more recent research has suggested that learned movements may be represented across both intrinsic and extrinsic reference frames, the latter including world- and object-based coordinate systems (Berniker et al., 2014, Brayanov et al., 2012, Leib and Franklin, 2025, Poh and Taylor, 2019). Leib and Franklin further proposed that the relative contribution of each coordinate system depends on its reliability, which is determined by the level of motor planning noise associated with that coordinate system (Leib and Franklin, 2025).

Importantly, intrinsic representations have typically been characterized using joint-angle changes (Fig. 1B). However, the muscle activation patterns required to produce a given joint-angle change depend on the limb configuration. Because muscle moment arms vary with joint configuration and muscle recruitment also depends on movement direction, movements aligned in joint space across different limb configurations may not be aligned in muscle space (Flanders and Soechting, 1990, Loeb et al., 1999). This raises the possibility that muscle-based coordinates (Fig. 1C) contribute to the intrinsic representation of learned movements in ways that are not captured by joint-based coordinates.

We first compared muscle activation patterns during reaching movements in two workspaces separated by a fixed shoulder rotation to determine whether movements aligned in joint space were also aligned in muscle space. We found that these movements were not necessarily aligned in muscle space and that the degree of misalignment varied systematically with movement direction. Based on these findings, we predicted that a muscle-space account, in which learned movements are represented using muscle-based coordinates, would predict different patterns of motor generalization following adaptation to clockwise (CW) and counterclockwise (CCW) visuomotor rotations. In contrast, a joint-space account, based on joint-based coordinates, would predict identical generalization patterns. We tested these predictions in Experiment 2 using 30° visuomotor rotations. Consistent with the muscle-space account, the results revealed a systematic shift between the generalization patterns, suggesting that muscle-based coordinates contribute to the representation of learned movements in ways not captured by joint-based coordinates alone. Together, these findings highlight the importance of considering the biomechanical relationship between joint and muscle coordinates when interpreting intrinsic representations of learned movements inferred from motor generalization paradigms.

## Methods

### Participants

Sixty-two healthy right-handed participants (20 females and 42 males; mean ± SD age: 25 ± 4 years) with normal or corrected-to-normal vision and no history of neurological disorders participated in this study after providing written informed consent. Thirty participants took part in Experiment 1, and 32 in Experiment 2, with the latter divided into two equal groups (n = 16 per group). The sample size in Experiment 1 (n = 30) was comparable to those used in previous studies examining muscle activity during reaching movements (Castillo-Escario et al., 2021, Dupuis et al., 2024). The sample size of Experiment 2 (n = 16 per group) was also consistent with that used in a previous visuomotor generalization study employing a similar experimental paradigm and was comparable to those used in previous studies employing a similar motor generalization paradigm (Poh and Taylor, 2019, Bao et al., 2022).

All experimental procedures were approved by the NICT Ethics Committee (N230122402) and were conducted in accordance with the Declaration of Helsinki and applicable national regulations.

### Apparatus

Both experiments were performed using the KINARM Exoskeleton Classic lab (BKIN Technologies, Kingston, ON, Canada), a dual-joint robotic apparatus that allows movements of shoulder and elbow. Participants were seated comfortably with their foreheads resting against the frame of the display and performed right-handed reaching movements while grasping a custom-designed handle. The experimental scene was displayed on a 47-inch digital monitor (resolution: 1920 × 1080 pixels; refresh rate: 60 Hz) and viewed through a semi-transparent mirror. During the experiment, participants could not see their hand, which was occluded by a horizontal physical shield beneath the mirror.

In Experiment 1, electromyographic (EMG) activity was recorded at 1000 Hz using surface electrodes (DELSYS, Natick, MA, USA) from nine muscles: (1) brachioradialis; (2) biceps brachii, short head; (3) biceps brachii, long head; (4) triceps brachii, lateral head; (5) triceps brachii, long head; (6) anterior deltoid; (7) middle deltoid; (8) posterior deltoid; and (9) pectoralis major. These nine muscles were selected to cover the main shoulder and elbow flexors and extensors, including biarticular muscles, acting during planar reaching. The experiments were controlled using the built-in BKIN software, Dexterit-E (version 3.10), which works with MATLAB Simulink Real-Time (version R2019a). Joint positions and velocities as well as cursor positions were sampled at 4000 Hz and stored at 1000 Hz. No EMG recordings were performed in Experiment 2.

### Motor task

In both experiments, participants performed a shooting reaching task. At the start of each trial, the participant’s hand, represented by a white dot (0.5 cm radius), was passively moved by the robot to a starting position represented by a grey circle (1 cm radius). Participants were required to maintain the cursor within the starting circle for a randomly selected duration between 500 and 800 ms. The starting circle then disappeared, and a target appeared as a grey circle (1 cm radius) located 10 cm (Experiment 1) or 7 cm (Experiment 2) from the start position. A 100-ms auditory cue signaled the preparation period. Four hundred milliseconds after the end of this cue, a second auditory cue of the same frequency instructed participants to initiate a reaching movement. Participants were instructed to move through the target without stopping. If participants initiated the movement before the onset of the second auditory cue, a message stating “Wait for the second beep!” was displayed on the monitor. The robot then stopped the movement, returned the participant’s hand to the start position, and the trial was repeated at the end of the block. From the onset of the second auditory cue, participants had 1500 ms to perform a ballistic reaching movement toward the target without corrective submovements. The movement period ended when the participant reached a radial distance of 10 cm (Experiment 1) and 7 cm (Experiment 2) from the start position or when 1500 ms had elapsed. Once the cursor left the start position, it remained visible throughout the movement period. If the target was successfully hit, it changed color to orange, and a reward sound was presented. If the participant missed the target, no visual change occurred and an error sound was presented. Regardless of movement accuracy, the cursor position remained frozen for 500 ms to provide endpoint error feedback. The participant’s hand was then returned to the start position by the robot.

To maintain similar movement speeds across participants, movement-time feedback was provided after each trial. Movement time was defined as the interval between movement onset (when the cursor velocity exceeded 5 cm/s) and the instant the cursor reached a radial distance of 10 cm (Experiment 1) or 7 cm (Experiment 2) from the start position, regardless of target accuracy. If movement time exceeded 250 ms, the message “too slow!” was displayed, whereas if movement time was shorter than 140 ms, the message “too fast!” was displayed. No message was displayed for movement times within the 140–250 ms range. Trials were not repeated based on movement-time feedback. To obtain clearer EMG signals, participants in Experiment 1 performed longer reaching movements than in Experiment 2 under the same movement-time criterion, resulting in higher movement speeds.

The task was performed in two workspaces: the frontal workspace, which served as the reference workspace in Experiment 1 (Fig. 2A) and the training workspace in Experiment 2 (Fig. 2B), and the lateral workspace, which served as the probe workspace in Experiment 1 (Fig. 2A) and the generalization workspace in Experiment 2 (Fig. 2B). In the frontal workspace, the start position was aligned with each participant’s body midline, with the elbow angle fixed at 90° and the shoulder angle at 62.5 ± 2.0° (mean ± SD across participants from both experiments). In the lateral workspace, the start position was rotated by 45° CW about the shoulder joint relative to the frontal workspace (Fig. 2A, B).

**Figure 2:**
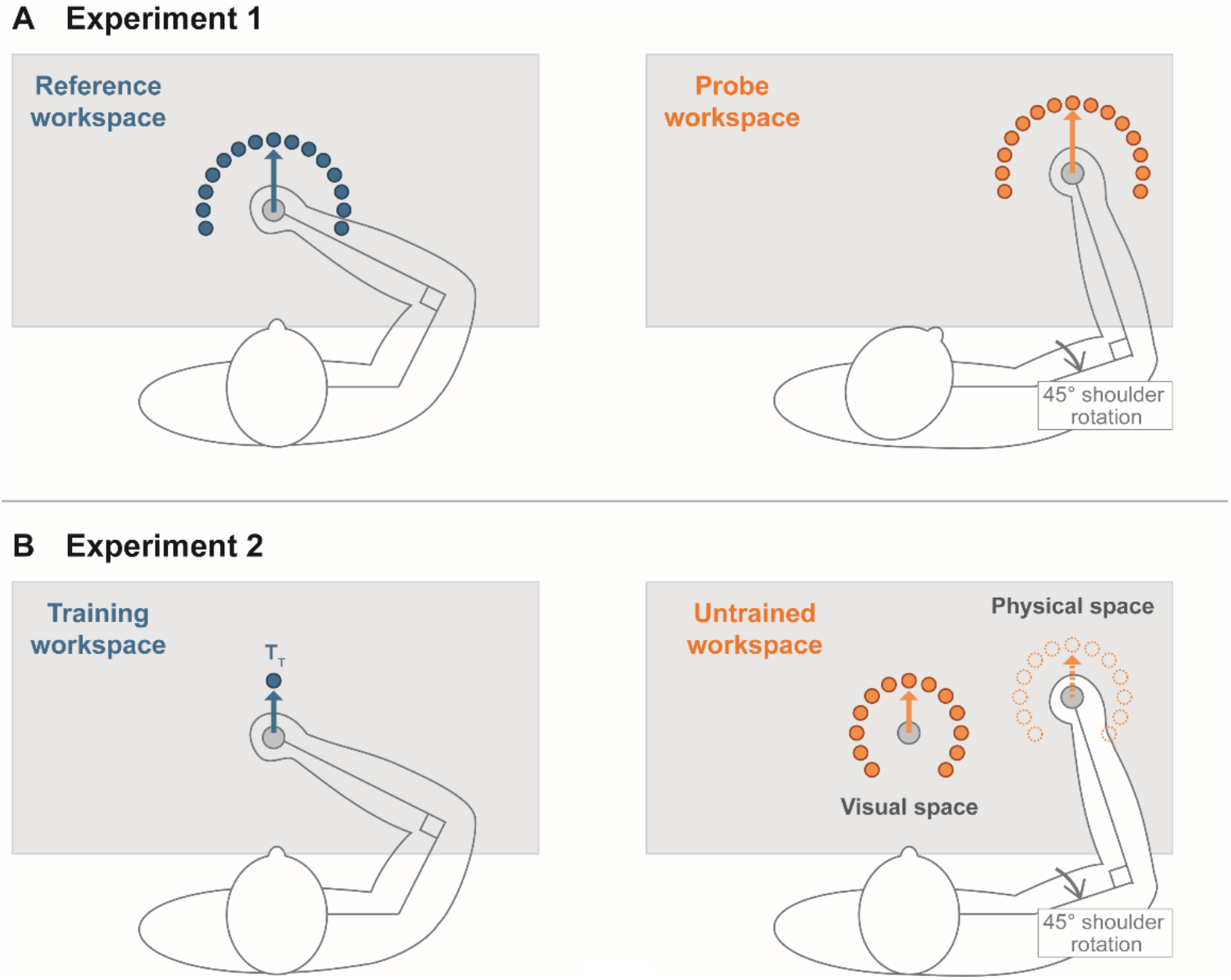
Experimental workspaces used in Experiments 1 and 2. (A) Experiment 1. Shooting reaching movements were performed in the reference and probe workspaces. The probe workspace was set by rotating the arm 45° CW about the shoulder joint relative to the reference workspace. Targets were arranged radially around the start position in both workspaces. (B) Experiment 2. Participants adapted to a visuomotor rotation in the training workspace and generalization was evaluated in the untrained workspace. During generalization measurements, the physical space, in which participants manipulated the handle, was dissociated from the visual space, in which the cursor, start position, and targets were displayed, such that gaze direction remained constant across workspaces.

### Procedures

#### Experiment 1

EMG activity was recorded from nine upper-limb muscles (see *Apparatus* section) while participants performed shooting reaching movements in the reference (frontal) and probe (lateral) workspaces (Fig. 2A). Thirty participants took part in the experiment. In both workspaces, 15 targets were arranged radially at 15° intervals, spanning from −105° to 105°, at a distance of 10 cm from the start position. The 0° direction corresponded to the forward reaching direction, with positive angles defined as CCW. The experiment consisted of four sets of 90 trials each (excluding repeated trials). The first and third sets were conducted in the reference workspace, whereas the second and fourth sets were conducted in the probe workspace. In each set, participants performed six reaching movements toward each of 15 targets in a randomized order. A break of at least 30 s was provided between sets. Before the experiment, all participants completed a familiarization session in which they performed one set in each workspace.

#### Experiment 2

We compared the generalization patterns following adaptation to 30° CW and 30° CCW visuomotor rotations to determine whether opposite rotations produced different generalization in the untrained workspace. Participants first learned a visuomotor rotation applied to the training target (T_T_) (left panel in Fig. 2B). A visuomotor rotation was defined as a rotation of the cursor movement direction relative to the hand movement direction. To minimize the potential effects of gaze direction on the generalization pattern, the physical space, in which participants manipulated the handle, was dissociated from the visual space, in which the cursor, start position, and targets were displayed (right panel in Fig. 2B) (Gurgone et al., 2025). There were two types of trials: training trials and generalization trials. During training trials, participants repeatedly reached toward the training target (T_T_ at 0° in the training workspace) with continuous cursor feedback to adapt to the visuomotor rotation. During generalization trials, participants reached toward one of the 13 generalization targets without cursor feedback to assess motor generalization while preventing feedback-based corrections and additional learning. The generalization targets were arranged radially at 22.5° intervals, spanning from −135° to 135° at a distance of 7 cm from the start position. Generalization trials were performed in either workspace, although the present analysis focused on the untrained workspace. During these trials, the target was displayed as a purple circle that did not change in size or color upon a target hit. In the untrained workspace, the visual stimuli were always displayed at the same location on the monitor as in the training workspace, keeping gaze direction constant across workspaces (Fig. 2B).

The experiment consisted of three sets: baseline, exposure, and generalization. The baseline set consisted of one training block followed by 13 generalization blocks. The training block included 20 trials to the training target (T_T_). In the generalization blocks, trials were organized into groups of seven, with three training trials followed by four generalization trials to one of the 13 generalization targets. Across the 13 generalization blocks, each generalization target was selected once in random order, resulting in four trials per target. During the baseline set, the visuomotor rotation was fixed at 0°. The subsequent exposure set consisted of 140 training trials, during which a visuomotor rotation was gradually introduced in increments of 3° every 10 trials following an initial 10 trials with no rotation. Thus, the rotation reached a maximum magnitude of 30° after 110 trials and was then maintained for the remaining 30 trials. Participants in one group adapted to a CW rotation, whereas those in the other group adapted to a CCW rotation. The generalization set followed the same trial structure as the baseline set, except that the visuomotor rotation was fixed at 30°. Short rest periods were provided between sets. Before the experiment, all participants completed a familiarization session in which they performed the task with and without visual feedback in both workspaces, with no visuomotor rotation.

Two additional experimental sets, unrelated to the main question of the present study, were conducted to evaluate generalization within the training workspace. These consisted of an additional baseline set and an additional generalization set performed in the training workspace before or after the corresponding untrained-workspace sets. The order of the training- and untrained-workspace sets was counterbalanced within each rotation group. These additional sets were excluded from the analyses presented here.

### Data analysis

Data handling and analysis for both experiments were performed using custom-written code in MATLAB R2017b (MathWorks). Participant characteristics are presented as mean ± SD and graphical data as mean ± SEM.

### Reaching behavior

Reaching behavior was quantified as angular deviation, defined as the angular difference between the hand movement direction and the target direction. The hand movement direction was defined by the vector from the hand position at movement onset to the hand position at peak velocity. Angular deviation was positive for CCW deviations and negative for CW deviations.

### EMG processing (Experiment 1)

For each trial and each of the nine recorded muscles, EMG activity was extracted from two trial phases: preparation and movement. Representative raw EMG recordings during reaching movements are provided in the Supplemental Material. The preparation phase began at 500 ms before movement onset and ended at movement onset, which was defined as 50 ms before the time point at which movement velocity exceeded 5 cm/s to account for the biomechanical delay between muscle activation and movement generation. The movement phase began at movement onset and ended when the cursor had moved 10 cm from the start position. EMG signals from both phases were full-wave rectified and filtered using a fourth-order low-pass Butterworth filter with a cut-off frequency of 20 Hz. To isolate movement-related EMG activity, the average EMG activity during the preparation phase was subtracted from the EMG signal during the movement phase. The resulting corrected EMG signal was then averaged across the movement phase to obtain a single EMG activity value for each trial. For each muscle, data points deviating by more than 5 SDs from the mean activation across all trials in an experimental set, in either the preparation or movement phases, were considered outliers and discarded. Additionally, trials with an absolute angular deviation greater than 15° were excluded across all muscles. The average number of discarded trials for each muscle was 3 ± 1 (mean ± SD). EMG activities were normalized to the maximum EMG value observed for each muscle after outlier exclusion. The normalized EMG data were then averaged across trials for each target, participant and experimental set.

### Muscle-pattern similarity analysis across workspaces (Experiment 1)

To assess whether movements aligned in joint space were also aligned in muscle space, we calculated the similarity between muscle activation patterns (hereafter referred to as muscle-pattern similarity) in the reference and probe workspaces (Fig. 3A, B). Cosine similarity has been used in previous studies to quantify the similarity between vectors representing muscle activity or muscle synergies (Hagio and Kouzaki, 2014, Nishida et al., 2017, Borzelli et al., 2018). In the current study, muscle activation patterns were represented as nine-dimensional vectors, with each dimension corresponding to the activation of one recorded muscle. Muscle-pattern similarity was then calculated as the cosine similarity between the muscle activation vector for a reference target and the muscle activation vector for a probe target (Fig. 3B). Similarity values were obtained for all combinations of reference and probe targets, yielding a similarity curve for each reference target expressed as a function of probe target direction (see Fig. 4A for an example). The peak of each similarity curve indicates the probe target direction most closely aligned with the corresponding reference target in muscle space. If joint and muscle spaces were aligned, similarity would peak at a probe direction shifted by −45° relative to the reference direction, matching the shoulder rotation (Fig. 2A).

**Figure 3:**
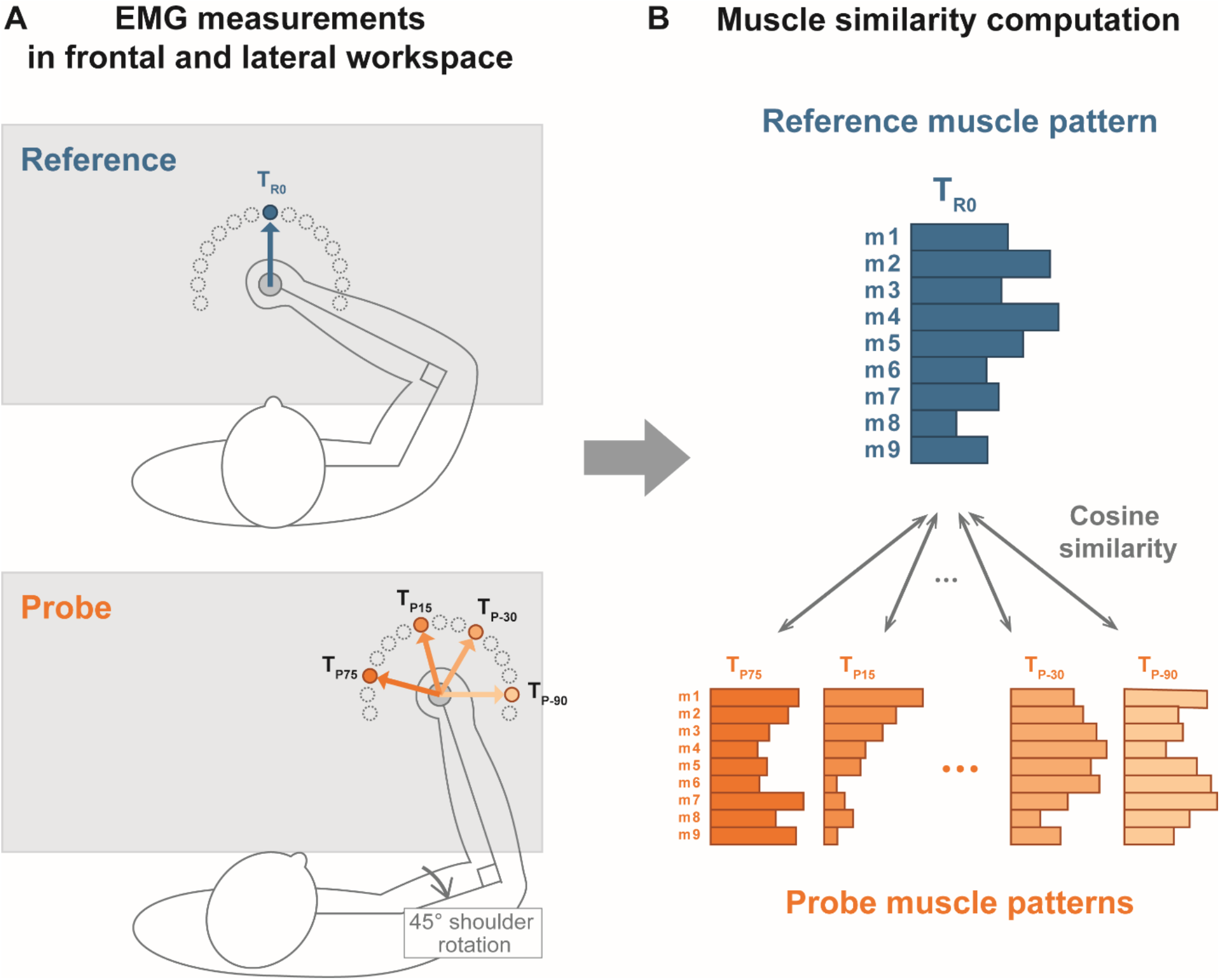
Calculation of muscle-pattern similarity across workspaces in Experiment 1. (A) EMG activity was recorded while participants performed shooting movements in the frontal (reference) and lateral (probe) workspaces. The two workspaces were separated by a −45° shoulder rotation. As an example, the 0° reference target and four representative probe targets are illustrated. (B) For each reference target, a nine-dimensional muscle activation vector, with each component representing the activity of one recorded muscle, was compared with the muscle activation vectors of all probe targets in the lateral workspace using cosine similarity. The bar lengths schematically represent the relative activation levels of individual muscles.

**Figure 4:**
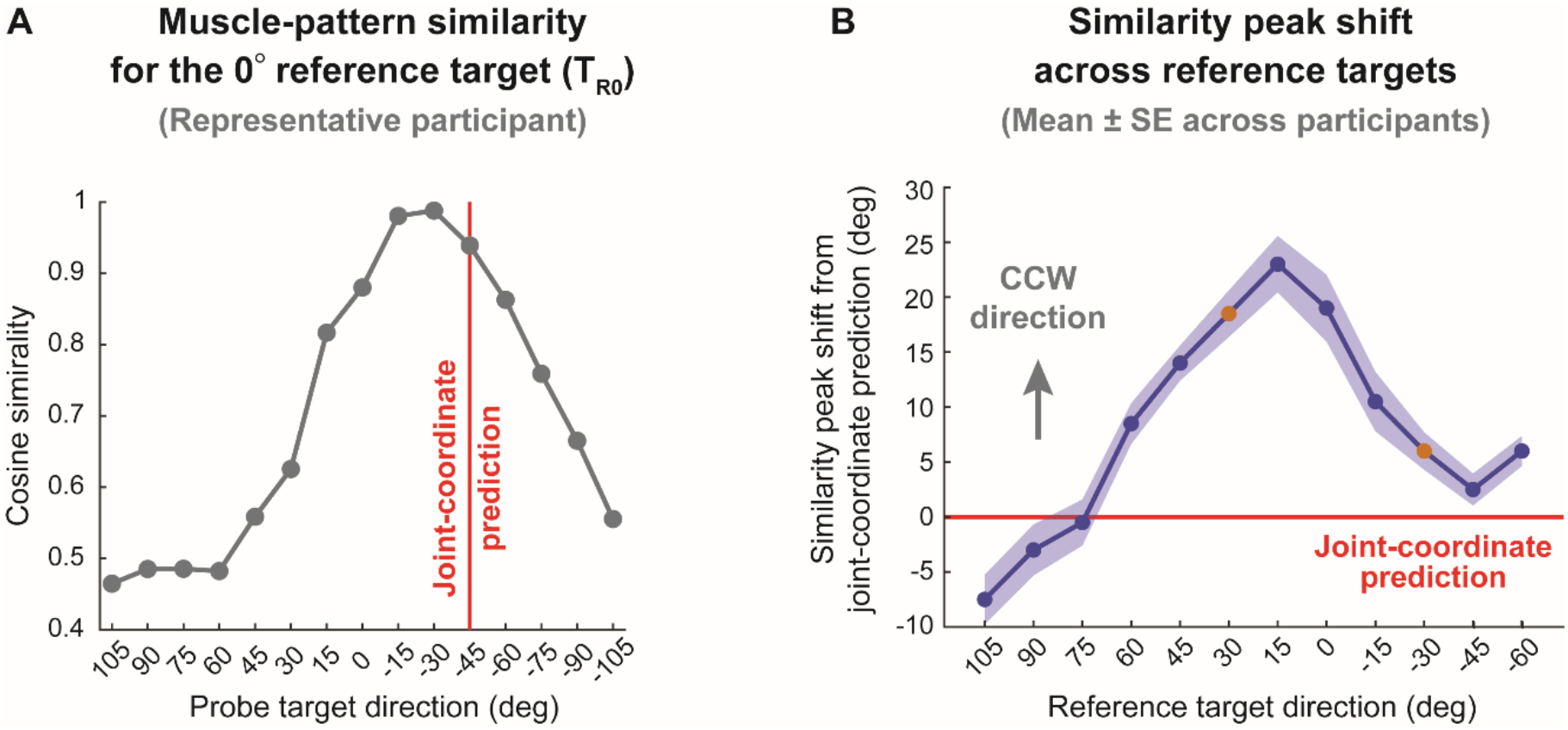
Muscle-pattern similarity and direction-dependent shifts from the joint-coordinate prediction in Experiment 1. (A) Muscle-pattern similarity curve for the 0° reference target from a representative participant. The peak (−30°) indicates the probe target direction with the highest muscle-pattern similarity, whereas the red vertical line indicates the joint-coordinate prediction (−45°). In this case, the similarity peak shift from the joint-coordinate prediction was 15°. (B) Similarity peak shifts relative to the joint-coordinate prediction plotted as a function of reference target direction. For each reference target, the shift was computed as the angular difference between the similarity peak and the corresponding predicted joint-coordinate direction (reference target direction minus 45°). Positive values indicate the shifts in the CCW direction relative to the joint-coordinate prediction. Shaded areas indicate ± SE across participants. The +30° and −30° reference directions used to derive the predictions for Experiment 2 are highlighted in orange.

This procedure yielded 15 muscle-pattern similarity curves per participant, one for each reference target. For each curve, the probe target direction corresponding to the maximum similarity was identified as the peak location. Because the theoretical joint-space prediction for the most peripheral reference targets (−75°, −90°, and −105°) fell outside the range of probe targets, peak-location analyses were restricted to reference targets from 105° to −60°. The resulting peak locations were analyzed using a one-way repeated-measures ANOVA (rANOVA) with reference target direction as the within-subject factor to assess whether peak location varied across reference targets. When the main effect was significant, Bonferroni-corrected one-sample t-tests were performed to compare the peak location for each reference target with the joint-space prediction.

### The generalization analysis of visuomotor rotation learning (Experiment 2)

In Experiment 2, we compared the generalization patterns in the untrained workspace following adaptation to the CW and CCW visuomotor rotations. Thirty-two participants were assigned to one of two groups: the CW rotation group (CW group) and the CCW rotation group (CCW group). To obtain a generalization pattern for each group, outlier trials were first excluded separately for each group, experimental set, and target by pooling data across participants. The mean angular deviation was calculated, and trials deviating by more than 3 SD from the mean were excluded. The angular deviations were then baseline-corrected by subtracting the mean angular deviation obtained in the baseline set for the corresponding target. The corrected angular deviations were averaged across trials for each participant and target. At this level, outliers were identified across participants separately for each group and target using MATLAB’s ‘isoutlier’ function with the default median-based criterion. As a result, 65 data points were excluded from the analysis (1% of group-by-participant-by-target data points). To present the learning effects of the two rotation groups using a common sign convention, the sign of the angular deviations for the CCW group was reversed before constructing the generalization curves and performing the statistical analyses. To compare the generalization patterns between the CW and CCW groups, a repeated-measures ANOVA (rANOVA) was performed, with group as a between-subject factor and target direction as a within-subject factor.

## Results

### Experiment 1: Misalignment between joint- and muscle-space across workspaces

Experiment 1 investigated whether movement directions aligned in joint space across workspaces are also aligned in muscle space. To address this question, we examined whether movement directions equivalent in joint space across workspaces were also associated with the highest muscle-pattern similarity. Muscle-pattern similarity was calculated between each reference target direction in the frontal workspace and all probe target directions in the lateral workspace. As a representative example, we first used the 0° reference target and calculated muscle-pattern similarity with all probe targets (Fig. 3A, B). The muscle-pattern similarity curve from a representative participant exhibited a bell-shaped tuning profile, with the peak indicating the probe target direction most closely aligned with the reference movement in muscle space. According to the joint-space account (see Methods), the predicted peak location corresponded to the reference target direction shifted by −45° (Fig. 2A). Thus, for the 0° reference target, the joint-space prediction was −45°. In the representative participant, however, the peak occurred at the −30° probe target rather than at the predicted −45° target (Fig. 4A).

We next examined whether this mismatch between joint space and muscle space was also observed across participants and across different reference movement directions. For each reference target, the probe target corresponding to the maximum muscle-pattern similarity was identified for each participant and compared with the corresponding theoretical joint-space prediction. The observed peak locations varied significantly across reference target directions (one-way rANOVA, F(11,319) = 21.51, p < 0.001, η_p_² = 0.43, Fig. 4). Importantly, Bonferroni-corrected one-sample comparisons revealed significant deviations from the joint-space predictions for all reference target directions except -45°, 75°, and 90° (uncorrected *p =* 0.10, 0.81, and 0.21, respectively). Together, these findings indicate that the correspondence between joint space and muscle space is movement-direction dependent. While joint and muscle spaces were closely aligned for some movement directions, substantial mismatches emerged for others.

### Behavioral predictions derived from Experiment 1

The movement-direction-dependent mismatch between joint space and muscle space observed in Experiment 1 (Fig. 4) provides distinct behavioral predictions for visuomotor generalization from the trained (frontal) to the untrained (lateral) workspace (Fig. 2B). Considering adaptation to visuomotor rotations of the same magnitude but opposite directions (CW and CCW), the joint-space account predicts identical generalization patterns for the two adaptations, whereas the muscle-space account predicts different generalization patterns (Fig. 5).

**Figure 5:**
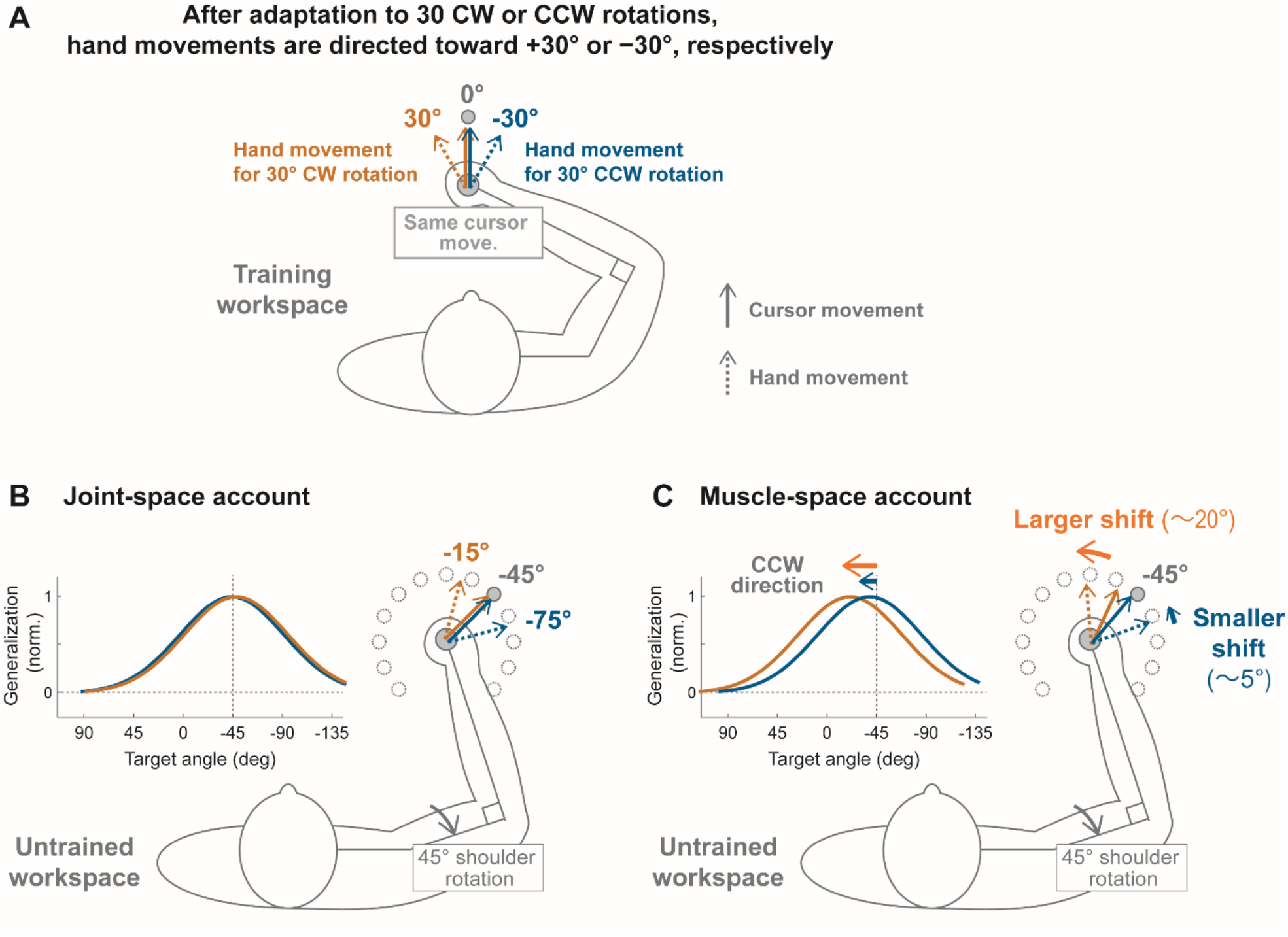
Schematic illustration of joint-space and muscle-space accounts for motor generalization following opposing visuomotor rotations. (A) Following complete adaptation to a 30° CW or CCW visuomotor rotation at the 0° training target, the adapted hand movements are directed toward +30° (orange) or −30° (blue), respectively, while the cursor moves toward the same target. (B) According to the joint-space account, the movement directions aligned with these adapted movements in the untrained workspace are −15° and −75°, respectively, corresponding to a −45° shift in joint-based coordinates. Thus, both rotations predict identical generalization patterns centered at the −45° target. (C) According to the muscle-space account, the direction-dependent mismatch observed in Experiment 1 produces different muscle-similarity peak shifts relative to the joint-coordinate predictions for the two adapted movements (approximately 20° for CW and 5° for CCW adaptation). Consequently, the generalization pattern following CW adaptation is predicted to shift further toward the CCW direction than that following CCW adaptation. The curves in (B) and (C) schematically illustrate the predictions of the two accounts rather than quantitative predictions of the generalization functions.

To illustrate these predictions, we considered 30° visuomotor rotations, a commonly used perturbation magnitude in visuomotor adaptation studies (Krakauer et al., 2000, Wang and Sainburg, 2005, Ikegami et al., 2010). Assuming complete adaptation to the 30° CW and 30° CCW rotations at the 0° training target and complete transfer of learning to the untrained workspace, adapted hand movements would be directed toward 30° and −30°, respectively (Fig. 5A). Under a joint-space account, which assumes that learned movements are represented exclusively in joint-based coordinates (Fig. 5B), the movement directions most closely aligned with these adapted movements in the untrained workspace are −15° and −75°, respectively. Because these movement directions are shifted by −45° relative to the adapted hand movements, both CW and CCW conditions predict maximal generalization to the −45° target in the untrained workspace. In contrast, a muscle-space account (derived from our muscle-pattern similarity analysis; Fig. 5C) yields different predictions. The +30° and −30° reference targets exhibited significantly different similarity peak shifts relative to the joint-coordinate predictions (18.5 ± 2.1° and 6.0 ± 1.7°, respectively; Bonferroni-corrected post hoc test, *p* = 0.002; Fig. 4B). Thus, the muscle-space account predicts that the behavioral generalization pattern following adaptation to a 30° CW visuomotor rotation should be shifted further in the CCW direction than that following adaptation to a 30° CCW visuomotor rotation. This prediction was tested in Experiment 2.

### Experiment 2: Behavioral test of the muscle-space account in visuomotor rotation generalization

Experiment 2 was performed to test the prediction derived from the muscle-space account: visuomotor adaptation to CW and CCW rotations should produce different generalization patterns. Participants adapted to either a 30° CW or 30° CCW visuomotor rotation in the training (frontal) workspace, and generalization was subsequently assessed in the untrained (lateral) workspace (Fig. 2B). Because the training target was located at 0° in the trained workspace, an extrinsic representation (Fig. 1A) predicts maximal generalization at the corresponding 0° target in the untrained workspace (Krakauer et al., 2000). Therefore, differences between the CW and CCW conditions were expected to appear primarily as shifts of the overall generalization pattern rather than as changes in the peak location. Both groups exhibited maximal generalization at the 0° untrained target. However, the overall generalization patterns were shifted relative to each other (Fig. 6). A repeated-measures ANOVA with group (CW vs. CCW) as the between-subject factor and target direction as the within-subject factor revealed a significant interaction between the two factors (F(12, 360) = 5.11, p < 0.001, η_p_² = 0.15), indicating that the generalization patterns differed between the two rotation groups. Consistent with the prediction derived from the muscle-pattern similarity analysis in Experiment 1 (Fig. 6), the generalization pattern following CW adaptation was shifted further toward the CCW direction than that following CCW adaptation.

**Figure 6:**
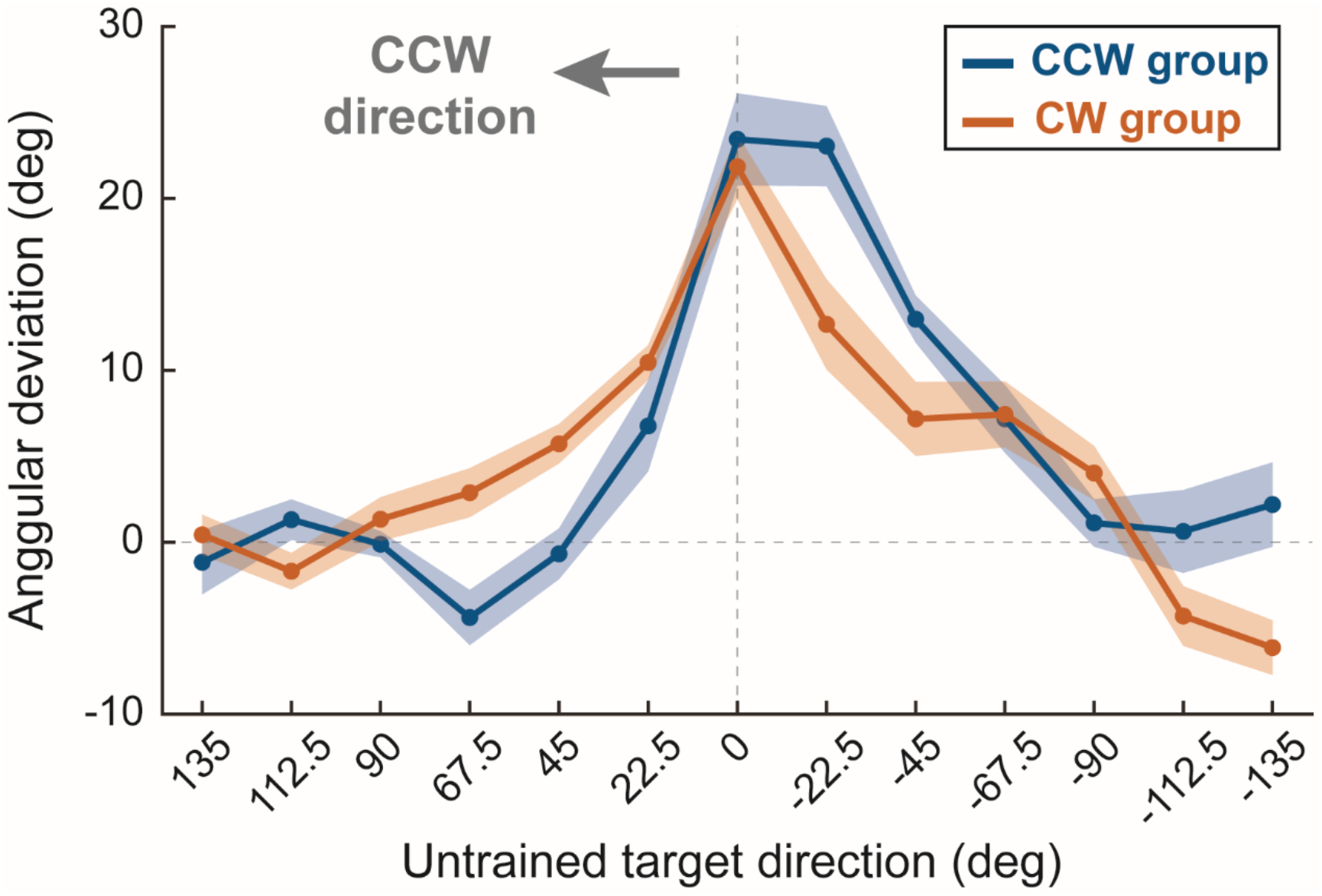
Generalization patterns following CW and CCW visuomotor rotation learning in Experiment 2. Mean baseline-corrected angular deviations (mean ± SE) are plotted as a function of untrained target direction. Because CW and CCW adaptation produced opposite signs of angular deviation, the sign of the CCW group’s angular deviations was reversed to allow direct comparison of the generalization patterns. Although both groups exhibited maximal generalization at the 0° untrained target, the overall generalization pattern for the CW group was shifted further toward the CCW direction than that for the CCW group, consistent with the prediction derived from the muscle-pattern similarity analysis in Experiment 1.

## Discussion

The present study examined whether movement directions aligned in joint space across different workspaces are also aligned in muscle space and, if not, whether such a mismatch can predict specific patterns of motor generalization. We found three main results. First, movement directions aligned in joint space across training and untrained workspaces were not always aligned in muscle space. Second, the degree of misalignment varied across reference target directions in the training workspace, indicating that the correspondence between joint and muscle coordinates is movement-direction dependent. Third, this mismatch led to distinct predictions for motor generalization following CW and CCW visuomotor adaptation, which was confirmed in Experiment 2. Together, these findings suggest that intrinsic representations of learned movements cannot be fully captured by joint-based coordinates alone and likely include a contribution from muscle-based coordinates.

### Why do joint- and muscle-space representations dissociate?

The mismatch between joint-space and muscle-space alignment observed in Experiment 1 is not unexpected from a biomechanical perspective. Joint-space accounts assume that movements producing the same pattern of joint-angle changes should be considered equivalent, regardless of the initial limb configuration. However, the mapping between joint-angle changes and muscle activation is highly dependent on posture. In the present study, the frontal and lateral workspaces differed in their initial shoulder configuration by 45°, resulting in different muscle lengths and mechanical conditions at movement onset. Because muscle moment arms vary with joint configuration, the same pattern of joint-angle changes can require different patterns of muscle activation depending on the workspace (Flanders and Soechting, 1990, Hik and Ackland, 2019). Consequently, movements aligned in joint space are not necessarily aligned in muscle space.

In addition to posture-dependent changes in muscle mechanics, muscle redundancy may further contribute to the observed dissociation. Distinct muscle activation patterns can generate similar joint-angle changes, particularly in multi-joint movements involving both monoarticular and biarticular muscles (Loeb et al., 1999, Prilutsky, 2000). Biarticular muscles are of particular interest because they coordinate movement across multiple joints and enable inter-joint mechanical interactions that are not available through monoarticular muscles alone. Consequently, identical joint-angle changes can be generated by different combinations of muscle activations depending on movement direction and limb configuration. Consistent with this idea, previous studies have shown that muscle activation patterns vary with posture even when movements are directed toward the same visual target (Sergio and Ostry, 1995). Together, these biomechanical properties of the musculoskeletal system provide a plausible explanation for why movement directions aligned in joint space were not always aligned in muscle space.

### Why does the mismatch matter for interpreting generalization paradigms?

Although the mismatch between joint and muscle spaces can be anticipated from biomechanical considerations, its implications for motor learning have remained largely unexplored. Generalization studies have traditionally characterized intrinsic representations using joint-based coordinate models, without explicitly considering whether discrepancies between joint and muscle spaces could affect the expression of learned movements across workspaces (Shadmehr and Mussa-Ivaldi, 1994, Brayanov et al., 2012, Berniker et al., 2014, Poh and Taylor, 2019). The present findings challenge this assumption. The joint-space account predicts that CW and CCW visuomotor rotations of equal magnitude generalize identically across workspaces, because the joint-space relationship between the trained and untrained workspaces is the same for both rotations. However, Experiment 1 showed that movements aligned in joint space are not always aligned in muscle space. Based on this mismatch, muscle-pattern similarity predicted that the generalization pattern following CW adaptation would be shifted toward the CCW direction relative to that following CCW adaptation. Experiment 2 tested these contrasting predictions and revealed the shift predicted by muscle-pattern similarity. This result was inconsistent with the conventional joint-space account.

Importantly, although our muscle-pattern similarity analysis provides a quantitative description of the correspondence between muscle activation patterns across workspaces, it does not provide a quantitative prediction of motor generalization. Nor does this analysis imply that the brain represents movements in exactly the same muscle space defined by our analysis, or specify how motor commands are computed to express learned movements in a new workspace. Such computations may involve muscle synergies or other neural and biomechanical mechanisms that are not captured by our analysis (d’Avella et al., 2003, Bizzi et al., 2008, Hagio and Kouzaki, 2014). Nevertheless, the muscle-pattern similarity measured in Experiment 1 qualitatively predicted patterns of motor generalization in Experiment 2 that could not be predicted from conventional joint-based coordinates alone. These findings suggest that muscle-based coordinates provide a behaviorally relevant description of motor generalization, beyond conventional joint-based coordinates. Together, these findings indicate that the mismatch between joint- and muscle-based coordinates can influence behavioral estimates of motor representations derived from generalization paradigms. Consequently, interpretations of intrinsic representations based solely on joint-coordinate alignment should be made with caution.

### Implications for theories of intrinsic representations

Our findings do not imply that learned movements are represented exclusively in muscle-based coordinates. Rather, the behavioral results suggest that muscle-based coordinates capture information that is not captured by joint-coordinate models alone. In this sense, muscle-based coordinates may complement, rather than replace, joint-based coordinates in the representation of learned movements. These findings are broadly consistent with previous studies proposing that learned movements are represented across multiple reference frames rather than within a single reference frame (Brayanov et al., 2012, Berniker et al., 2014, Leib and Franklin, 2025). However, whereas previous work has mainly focused on the relative contributions of intrinsic and extrinsic reference frames, the present results suggest that intrinsic representations may also involve multiple body-based coordinates. Thus, the distinction between intrinsic and extrinsic reference frames alone may be insufficient to fully describe the representation of learned movements, as joint-based and muscle-based coordinates may both contribute to intrinsic representations.

Consistent with this possibility, neurophysiological studies have identified motor cortical neurons encoding movement-related information in extrinsic, joint-, and muscle-based coordinates (Kakei et al., 1999, 2001). Although we did not directly examine neural activity, our findings open new avenues for investigating how neural circuits represent information across multiple reference frames to support motor learning and generalization.

## Conclusion

The present study demonstrated that movement directions aligned in joint space are not always aligned in muscle space and that this mismatch has important consequences for motor generalization. By combining muscle-pattern similarity analysis and behavioral testing, we showed that muscle-space account predicts differences in motor generalization that are not predicted by the conventional joint-space account. These findings suggest that muscle-based coordinates provide distinct information relevant to the intrinsic representation of learned movements and further highlight the importance of considering underlying biomechanics when interpreting motor representations inferred from generalization paradigms.

## Supporting information

Supplemental Table 1: Representative raw EMG recordings during reaching movements in Experiment 1.

## Acknowledgments

We thank Noriko Karasudani, Mayura Fujita, Mayumi Irikawa for their help in conducting the experiments and the members of the CiNet Motor Control Unit for their helpful comments.

