## Supplemental Table 1: Representative raw EMG recordings during reaching movements in Experiment 1. for "Direction-dependent misalignment between joint and muscle spaces predicts different generalization patterns following adaptation to opposing visuomotor rotations"

### Supplemental Material

#### Supplemental figure

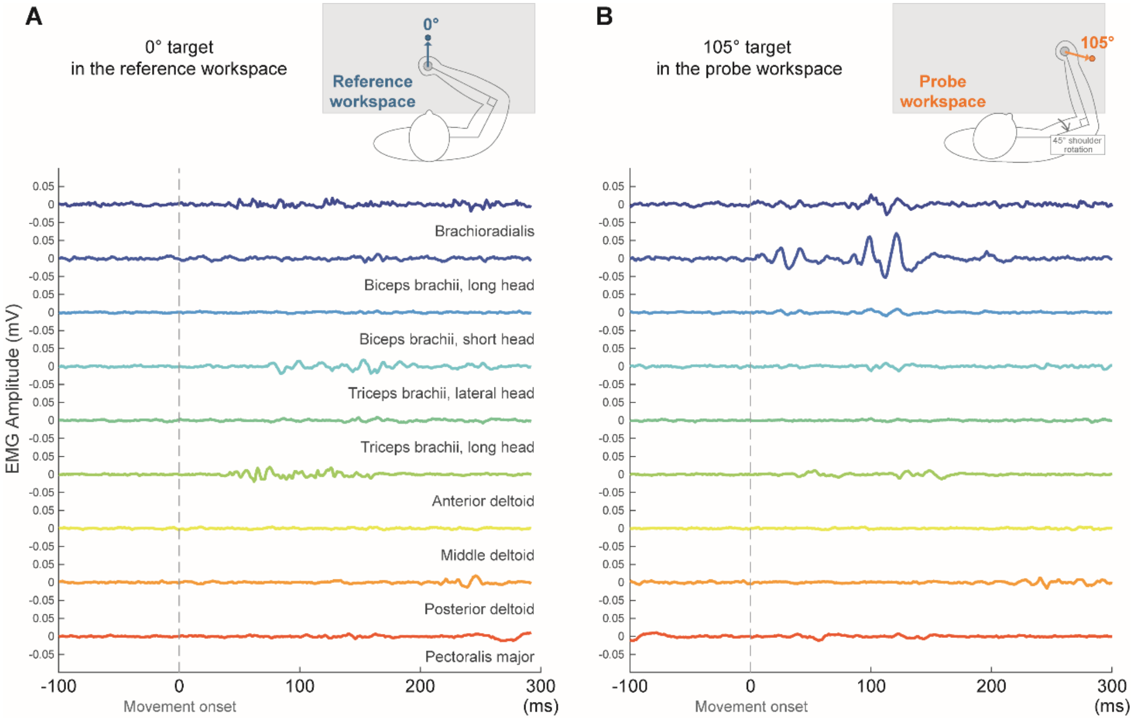


***Fig. S1. Representative raw EMG recordings during reaching movements in Experiment 1.*** *Single-trial raw EMG signals from nine upper-limb muscles during reaching toward (A) the 0° target in the reference workspace and (B) the 105° target in the probe workspace. The upper illustrations show the corresponding arm configurations and target directions. Dashed vertical lines indicate movement onset (0 ms). EMG traces are vertically offset for clarity.*
